# The effect of inclusion criteria on the functional properties reported in mouse visual cortex

**DOI:** 10.1101/2020.05.12.091595

**Authors:** Natalia Mesa, Jack Waters, Saskia E. J. de Vries

## Abstract

Neurophysiology studies require the use of inclusion criteria to identify neurons responsive to the experimental stimuli. Five recent studies used calcium imaging to measure the preferred tuning properties of layer 2/3 pyramidal neurons in mouse visual areas. These five studies employed different inclusion criteria and report different, sometimes conflicting results. Here, we examine how different inclusion criteria can impact reported tuning properties, modifying inclusion criteria to select different sub-populations from the same dataset of almost 10,000 layer 2/3 neurons from the Allen Brain Observatory. The choice of inclusion criteria greatly affected the mean tuning properties of the resulting sub-populations; indeed, the differences in mean tuning due to inclusion criteria were often of comparable magnitude to the differences between studies. In particular, the mean preferred temporal frequencies of visual areas changed markedly with inclusion criteria, such that the rank ordering of visual areas based on their temporal frequency preferences changed with the percentage of neurons included. It has been suggested that differences in temporal frequency tuning support a hierarchy of mouse visual areas. These results demonstrate that our understanding of the functional organization of the mouse visual cortex obtained from previous experiments critically depends on the inclusion criteria used.

## INTRODUCTION

Five recent studies have employed 2-photon calcium imaging to compare spatial frequency (SF) tuning, temporal frequency (TF) tuning, orientation selectivity, and directional selectivity of neurons across mouse visual cortical areas (**Table 1, Figure 1**) (Andermann et al. 2011; Marshel, Garrett et al. 2011; Roth, Helmchen, and Kampa 2012; Tohmi et al. 2014; Sun et al. 2016). Some results were consistent across studies, e.g. the mean preferred TF of neurons in area AL was greater than those in V1 (**Figure 1A**), but there were also differences between studies, e.g. some studies found that the mean preferred TF of neurons in PM was greater than those in V1 while others found the opposite. Further, the magnitudes of average TF tuning, orientation selectivity index (OSI), and direction selectivity index (DSI) in individual visual areas as well as the rank order of these properties between visual areas differed across studies (**Figure 1**). All five studies imaged layer 2/3 of mouse visual cortex and activity was evoked with a drifting grating stimulus, but the studies differed in anesthesia state, calcium indicator, and in the inclusion criteria used in analysis (**Table 1, columns 2-4**). It is likely that all these differences contribute to the contrasting results. Here, we leverage a single large and open dataset, the Allen Brain Observatory, to quantify the impact of the choice of inclusion criteria on the measurement of tuning properties of neurons in mouse visual areas.

**Table 1.**
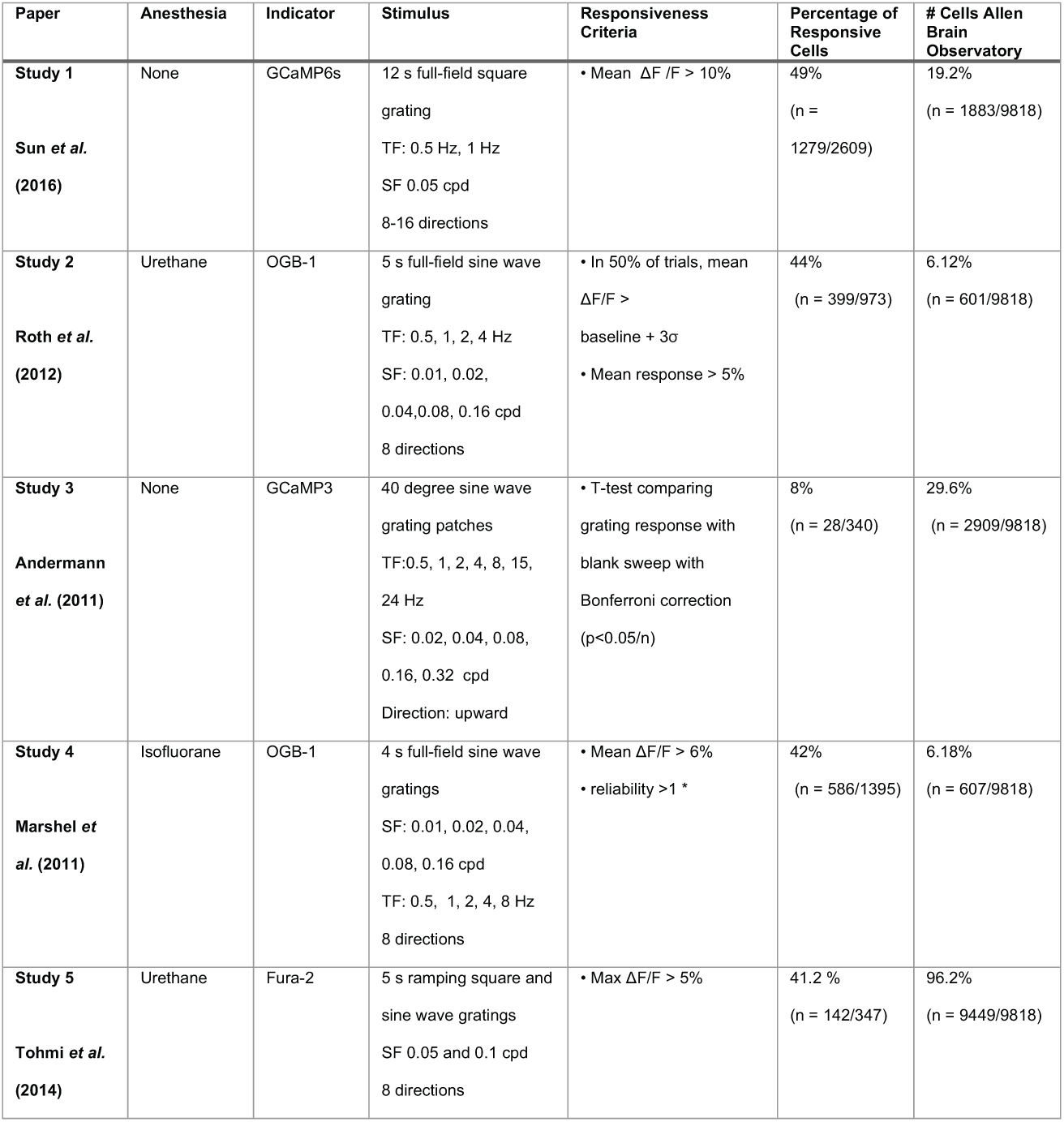
Summary of the experimental conditions and inclusion criteria used in published studies.

**Figure 1.**
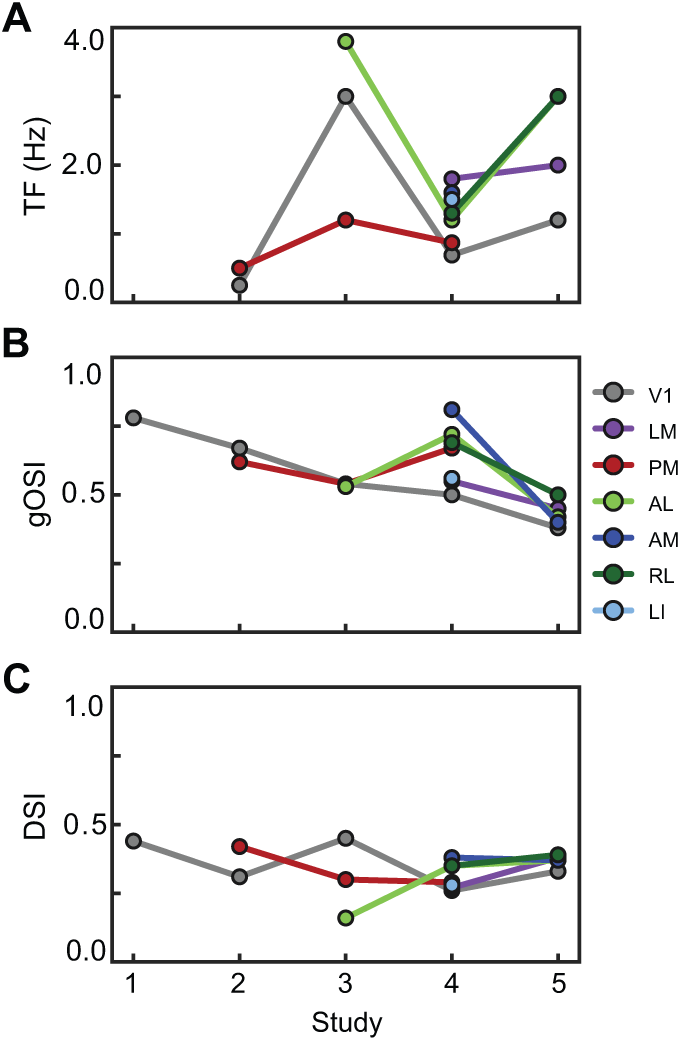
Tuning characteristics in published studies. (A) Mean temporal frequency (TF) tuning of seven visual areas reported in five published studies. (B - C) Same as in A, but reporting the orientation selectivity index (OSI) and the direction selectivity index (DSI).

Calcium imaging studies usually require the use of inclusion criteria to select neurons that are deemed to be “active” or “responsive” such that the derived analysis of their activity is relevant to the aims of the experiment and not a quantification of noise. As the measured fluorescence shows continuous fluctuations, these criteria serve to identify which fluctuations reflect signal rather than noise. Criteria are often based on the amplitude of the fluorescence change, e.g. a threshold on the mean or median change in fluorescence over multiple trials, or its reproducibility, e.g. a statistically significant stimulus-evoked change in fluorescence on a subset of trials. Naturally, some neurons exhibit large-amplitude changes in fluorescence on every trial in response to a preferred stimulus and fulfil both amplitude and reproducibility criteria (**Figure 2A-C**). Many neurons display reproducible, small-amplitude changes (**Figure 2D-F**) or large-amplitude changes in fluorescence on only some trials (**Figure 2G-I**). Although not often used as the basis for inclusion criteria, other features of the fluorescence traces, such as periodicity in the fluorescence in response to a periodic stimulus such as a drifting grating (**Figure 2I**) and tuning to stimulus characteristics such as orientation and temporal frequency (**Figure 2C, H, I**), may also be suggestive of stimulus-evoked activity (Neill & Stryker, 2008).

**Figure 2.**
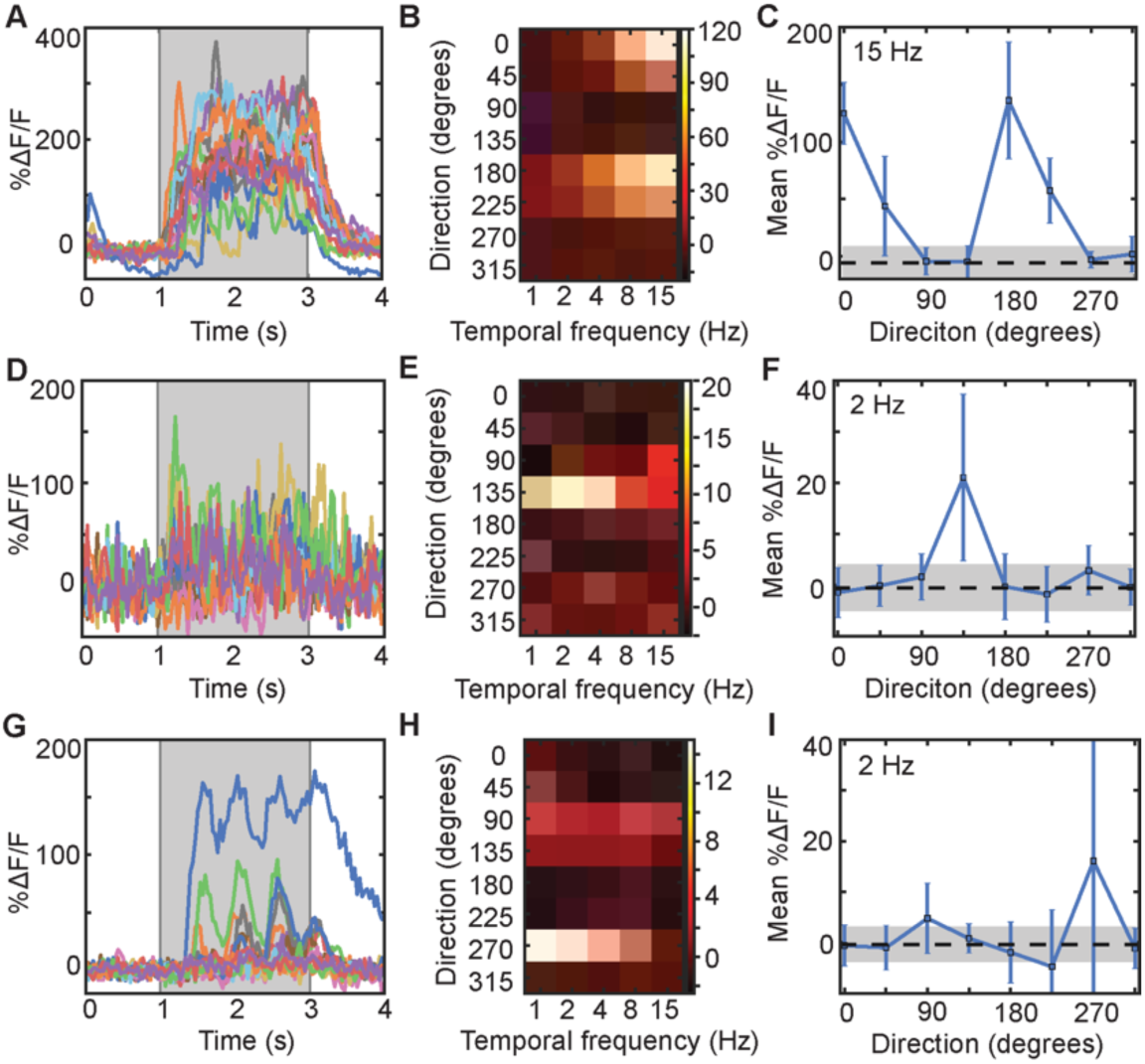
Example cells that pass different sets of inclusion criteria. (A) All dF/F responses to the preferred stimulus condition of a cell that passes all of the criteria compared here. (B) Heatmap of mean %ΔF/F responses to each stimulus condition (TF x orientation) for the same cell as in A. (C) Mean (± standard deviation) %ΔF/F responses to stimuli of different orientations for the same cell as in A. (D-F) Same as in A-C, but with a cell that passes most criteria, but not Study 2. (G-I) Same as in A-C, but with a cell that only passes Study 1 criteria.

Each of the five studies used different inclusion criteria and it is unclear whether these different criteria select the same or different neurons and how they impact the distribution of measured responses to visual stimuli across the population. Here we explore the effects of inclusion criteria on results from a single large dataset, eliminating the effects of different experimental conditions. We used recordings from the Allen Brain Observatory, a database of physiological activity in visual cortex measured with 2-photon calcium imaging from adult GCaMP6f transgenic mice (de Vries, Lecoq, Buice et al., 2020). We found that tuning properties varied with inclusion criteria, in some cases changing the rank order of tuning properties across mouse cortical visual areas.

## RESULTS

The five studies employed a range of inclusion criteria, selecting 8-49% of the neurons in their respective studies (Table 1). We applied the five different inclusion criteria to the Allen Brain Observatory, a large 2-photon calcium imaging data set. We restricted our analysis to layer 2/3 excitatory neurons imaged 175-250 μm below the pia in Cux2-CreERT2;Camk2a;Ai93 and Slc17a7-IRES2-Cre;Camk2a;Ai93 mice, yielding a dataset of fluorescence recordings from 9,818 neurons. The inclusion criteria from the 5 studies were all based on one or both of the amplitude and the trial-to-trial variability of the evoked responses and we therefore calculated the mean and standard deviation of the response of each neuron to its peak stimulus condition (the direction and TF that evoked the largest mean response). Different inclusion criteria selected different numbers of neurons (6-94% of 9,818 neurons, Table 1 Column 7) and when we visualize the neurons by plotting response mean vs standard deviation, these neurons occupy different but often overlapping locations (**Figure 3A, C**). The mean OSI and DSI values derived using these different criteria covered similar ranges to those in the published studies, consistent with the idea that effects of inclusion criteria might be sufficient to account for some of the disparate results across published studies (**Figure 3B**).

**Figure 3.**
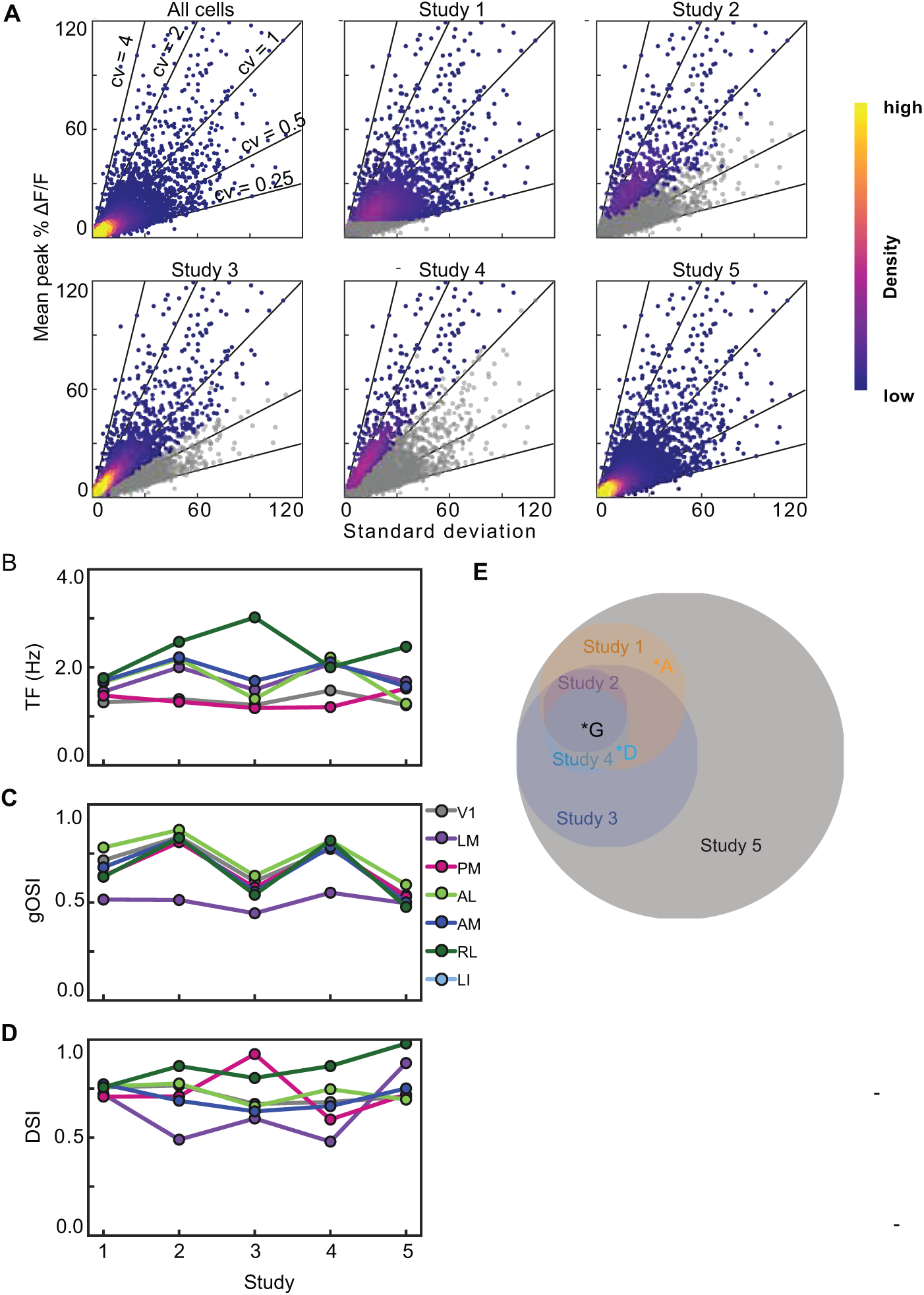
Most studies select for neurons along similar axes of the data. (A) Six density plots of the mean response at the preferred stimulus condition (% dF/F) against the standard deviation of the responses at the preferred stimulus condition where each point represents a single neuron. For each study, colored neurons are those selected for by inclusion criteria. Heatmap represents the density of neurons. (B-D) Tuning characteristics after inclusion criteria are applied to Allen Brain Observatory Dataset. B shows mean TF tuning of six visual areas when different inclusion criteria are applied. C and D show the mean OSI and DSI of six visual areas, respectively. (E) Venn Diagram of neurons that were selected for by each inclusion criteria. Area of circles represents the number of neurons. Example neurons from Figure 2 are indicated by letters.

Using the coefficient of variation (CV = standard deviation/mean) as a measure of the robustness of the response, we asked how increasing the number of neurons selected, from the most robust (lowest CV) to the least (highest CV), affects the computed tuning metrics. For some metrics, including more neurons affected tuning properties by almost as much as the differences between studies. For example, increasing included neurons changed the mean preferred TF for V1, PM, and AL and the rank order of these three areas, such that AL and PM display different mean TFs when only the top decile are included, but have the same mean TF when all neurons are included (**Figure 4A-D**). Within V1, the change in mean TF reflects the fact that the highest decile (10% with highest CV) shows a broader distribution of preferred TF than the lowest decile (**Figure 4B,C**). In contrast, the effect on OSI was negligible (**Figure 4E-H**). Finally, increasing the number of neurons included increased the mean DSI, and did so consistently across all visual areas (**Figure 4I-L**). The increase in DSI reflects the fact that many of the neurons in the lowest decile have a DSI of 1, whereas the neurons in the highest decile have a uniform distribution of DSIs (**Figure 4J,K**).

**Figure 4.**
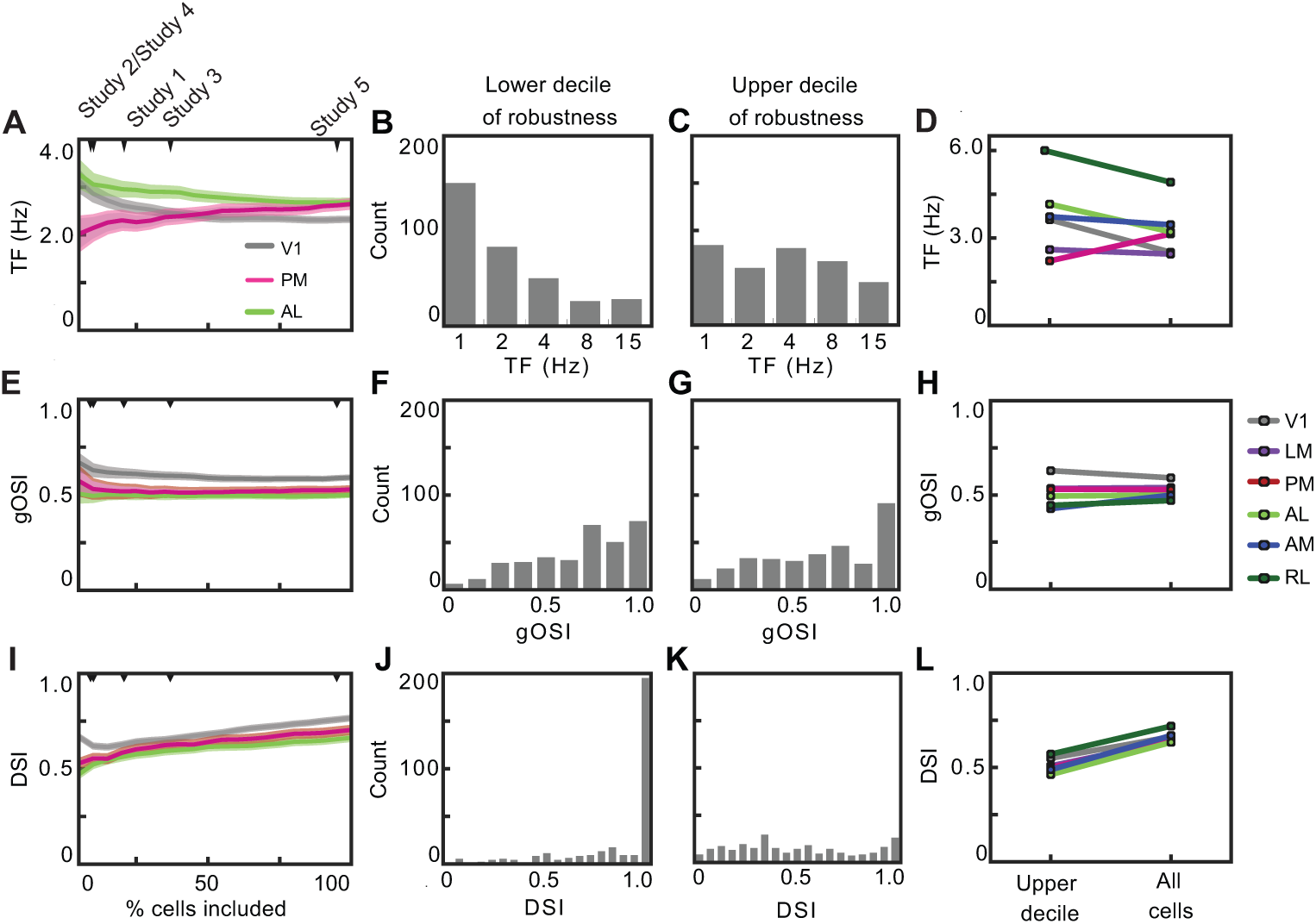
Tuning characteristics of neurons based on robustness. (A, E, I) Mean TF, OSI, and DSI tuning of neurons in V1, AL, and PM based on what percentage of most robust cells (cells with low coefficient of variantion) are included in the analysis. Shaded regions indicate SEM. (B) Histogram of TF tuning of 10% least robust cells. (C) Histogram of TF tuning of 10% most robust cells. (F, G, J, K) Same as in B, C but with OSI and DSI. (D, H, L) Mean TF, OSI, and DSI tuning of neurons in all visual areas in 10% most robust neurons versus the entire population of neurons.

## DISCUSSION

We applied different inclusion criteria to the Allen Brain Observatory 2-photon dataset to examine how these criteria impact the reported tuning properties across visual areas after experimental differences are eliminated. That different inclusion criteria selected different subsets of neurons might not be surprising, but the extent of the differences between selected neurons was substantial. One key difference was in the numbers of neurons selected. To examine how including more, or fewer, neurons could impact the tuning properties, we used CV as a metric of robustness and shifted our threshold for inclusion. Mean TF, OSI and DSI changed differently with the robustness of the responses of the underlying neurons. The preferred TF was the most sensitive, OSI the least sensitive.

Our results offer one possible explanation why published studies comparing TF, OSI and DSI across mouse visual areas have produced different results for TF and more similar results for OSI and DSI. Mean TF tuning is more sensitive than OSI and DSI to the neurons selected. As a result, comparison across studies is difficult and there remains considerable uncertainty in the mean TF and the rank order of TF tuning across mouse visual areas.

The lesser sensitivity of OSI and greater sensitivity of DSI and TF to the inclusion threshold may result, in part, from the lesser sensitivity to noise of the OSI than of the DSI and TF calculations. The neurons with the noisiest responses (greatest CV) commonly displayed DSI ∼1, which is inevitable when the response to the null direction is 0. The response to the preferred direction need not be large and could even result from a single trial having just a small amplitude fluorescence change. As the preferred TF is the TF at which the neuron has its largest response regardless of amplitude or reliability, the TF tuning is similarly sensitive to small numbers of noisy events. In contrast, OSI is calculated from the responses to all eight directions of drifting grating and is thus less sensitive to a small amplitude response in one condition. It is likely that the OSI measurement is more robust to noise than DSI and TF and this is why mean OSI across visual areas changes little with selected neurons.

We used CV to examine how including more neurons can impact the reported results, as one of the big differences of the criteria is the number of neurons they select from our dataset. But this is not the only difference between these criteria. This is evident from the Venn diagram that reveals the different criteria select somewhat non-overlapping groups of neurons and is not a set of concentric circles. This reveals that different inclusion criteria use features of the neural responses other than the size and reliability of neurons’ responses to their preferred condition. For instance, the statistical tests employed in Studies 3 and 4 also depend on the size and reliability of the neurons’ responses to the blank sweep. A result of this is that while OSI is not impacted by including more neurons based on CV, it is impacted by applying the different inclusion criteria from the studies (see **Figure 3C, 4E**). This reveals that additional dimensions in the response space can reveal different subsets of neurons.

Our results illustrate how inclusion criteria can play a role in determining the tuning properties visual areas. Inclusion criteria are unlikely to account for all of the differences observed between the original studies, indicating that other experimental factors are important. Other factors likely include anesthesia state, the type of anesthesia used, the calcium indicator, and image brightness. Brain state can modulate neural responses in visual cortex, and anesthesia in particular can impact both the spontaneous and evoked responses. The type of anesthesia can also be a factor, with urethane impacting spontaneous and evoked firing rates but not OSI (Niell and Stryker 2010) and atropine affecting OSI but not spontaneous firing rate, evoked firing rate, DSI, preferred TF, or preferred SF (Durand et al. 2016). Calcium indicators have different sensitivities and signal to noise properties (Hendel et al. 2008; Chen et al. 2013), such that thresholds in mean DF/F appropriate for one indicator might not be appropriate for another. Most of the inclusion criteria selected ∼40-50% of neurons when applied to the matched study, but when applied to the Allen Brain Observatory data the percentage of neuron included often differed substantially, presumably because experimental conditions such as indicator brightness differed across studies. For example, simple thresholds on peak DF/F cannot be applied uniformly across different calcium indicators. Thus it seems unlikely that a single set of inclusion criteria would be appropriate across a wide range of experimental conditions.

Functional specialization of the higher visual areas in mouse cortex has been interpreted as evidence of parallel streams (Andermann et al. 2011; Marshel, Garrett et al. 2011). For example, V1 is thought to transfer low TF, high SF information to PM, the putative gateway to the dorsomedial stream (Glickfeld et al. 2013; Polack and Contreras 2012; Lopez-Aranda et al. 2009). However, in some studies, neurons in V1 and PM have similar mean TF tuning (with PM’s being 1.3-2x that of V1) (Roth, Helmchen, and Kampa 2012; Marshel et al. 2011), while others show that mean TF tuning in PM neurons that is 1/3 that of V1 neurons (Andermann et al. 2011). Our results indicate that in the most robust neurons, V1 has a higher TF tuning than PM, but in the least robust neurons, PM has a higher TF tuning than V1, potentially explaining the discrepancies between studies. Since TF is sensitive enough to inclusion criteria to change the relative order of TF tuning, it is difficult to interpret the relative TF tuning between visual areas currently. The most appropriate inclusion criteria would take into account how downstream targets filter or weight inputs and how robustness factors into that weighting. Since we don’t know what this weighting is, we must be cautious in drawing conclusions about functional organization from these analyses.

## METHODS

### Stimulus and Dataset

We used calcium imaging recordings from the Allen Brain Observatory, a publicly available dataset that surveys physiological activity in the mouse visual cortex (de Vries, Lecoq, Buice *et al*., 2020). We specifically used the responses to the drifting grating stimulus in this dataset. This stimulus consisted of a 2s grating followed by a 1s mean luminance grey period. Six temporal frequencies (1, 2, 4, 8, 15 Hz), eight different directions, and one spatial frequency (0.04 cpd) were used. Each grating condition was presented 15 times.

Data analysis was performed in Python using the AllenSDK. The evoked response was defined as the mean dF/F during the 2s grating presentation. Responses to all 15 stimulus presentations were averaged together to calculate the mean evoked response.

We restricted our analysis to cells in layer 2/3 (175 um below pia, exclusive) of transgenics lines Cux2-CreERT2;Camk2a-tTa;Ai93 and Slc17a7-IRES2-Cre;Camk2a-tTa;Ai93, which express GCaMP6f in neural populations in layer 2/3 and throughout neocortex, respectively. A total of N = 9818 cells from 52 mice (28 male, 24 female) were used for this analysis.

### Metrics

The preferred direction and temporal frequency condition was defined as the grating condition that evoked the largest mean response. In order to compute the average TF tuning of a population of neurons, these TF values were first converted an octave scale (base 2), averaged, then converted back to a linear scale and reported.

Directional selectivity was computed for each neuron as

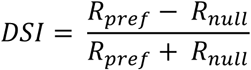

where *R*_*pref*_ is the mean response at to the preferred direction and *R*_*null*_ is the mean response to the opposite direction.

Orientation tuning was computed for each neuron using the global orientation selectivity index (OSI), (Ringach et al., 1997) defined as:

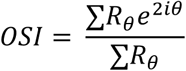

Where *R*_*θ*_is the mean response at each orientation *θ*.

The coefficient of variance (CV) was used as our metric to determine robustness. CV was calculated for each neuron as the ratio of standard deviation of the 15 responses to the preferred condition (mean dF/F over the 2s stimulus presentation) to the mean evoked response (see above). A low CV would indicate high robustness.

### Inclusion criteria

Published studies used the following inclusion criteria, which we applied to cells in the Allen Brain Observatory Dataset in the following manner:

Study 1: The mean evoked response (dF/F) to the preferred stimulus condition is greater than 10%. (Sun et al. 2015)

Study 2: In 50% of trials, the response is (1) larger than the 3x the standard deviation of the pre-stimulus baseline and (2) larger than 5% dF/F. (Roth, Helmchen, and Kampa 2012)

Study 3: Paired t-test (p > 0.05) with Bonferroni correction comparing the mean evoked response during the blank sweeps with mean evoked responses to preferred stimulus condition. (Andermann et al. 2011)

Study 4: (1) The mean response (dF/F) to any stimulus condition is is greater than 6%. And (2) reliability>1 where:

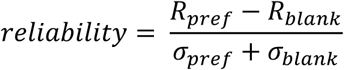

(Marshel, Garrett et al. 2011)

Study 5: The maximum fluorescence change (dF/F) during the 2s stimulus presentation block to any stimulus condition was greater that 4%. (Tohmi et al. 2014)

### Code accesibility

The code used for these analyses is available at https://github.com/nataliamv2/inclusion_criteria

## Notes

http://observatory.brain-map.org/visualcoding/

### Competing Interest Statement

The authors have declared no competing interest.

